# Specific Oxylipins Enhance Vertebrate Hematopoiesis via the Receptor GPR132

**DOI:** 10.1101/313403

**Authors:** Jamie L. Lahvic, Michelle Ammerman, Pulin Li, Megan C. Blair, Emma Stillman, Anne L. Robertson, Constantina Christodoulou, Julie R. Perlin, Song Yang, Nan Chiang, Paul C. Norris, Madeleine L. Daily, Shelby E. Redfield, Iris T. Chan, Mona Chatrizeh, Michael E. Chase, Olivia Weis, Yi Zhou, Charles N. Serhan, Leonard I Zon

## Abstract

Epoxyeicosatrienoic acids (EETs) are endogenous lipid signaling molecules with cardioprotective and vasodilatory actions. We recently showed that exogenous addition of 11,12-EET enhances hematopoietic induction and engraftment in mice and zebrafish. EETs are known to signal via a G-protein coupled receptor(s), and significant research supports the existence of a specific high-affinity receptor. Identification of a hematopoietic specific EET receptor would enable genetic interrogation of the EET signaling pathway and perhaps clinical use of this molecule. We developed a bioinformatic approach to identify the EET receptor based on the expression of GPCRs in cell lines with differential responses to EETs. We found 10 candidate EET receptors that are commonly expressed in three EET-responsive human cell lines, but not expressed in an EET-unresponsive line. Of these candidates, only GPR132 showed EET-responsiveness *in vitro* using a luminescence-based assay for β-arrestin recruitment. Knockdown of zebrafish *gpr132b* prevented EET-induced hematopoiesis, and marrow from GPR132 knockout mice showed decreased long-term engraftment capability. In contrast to the putative high-affinity EET receptor, GPR132 is reported to have affinity for additional fatty acids *in vitro,* and we found that these same fatty acids enhance hematopoietic stem cell specification in the zebrafish. We conducted structure-activity relationship analyses using both *in vitro* and *in vivo* assays on diverse medium chain fatty acids. Certain oxygenated, unsaturated free fatty acids showed high activation of GPR132, while unoxygenated or saturated fatty acids had lower activity. Absence of the carboxylic acid moiety prevented activity, suggesting that this moiety is required for receptor activation. GPR132 responds to a select panel of polyunsaturated, oxygenated fatty acids to enhance both embryonic and adult hematopoiesis.

## Introduction

Eicosanoids are endogenous bioactive mediators derived from arachidonic acid, responsible for a variety of physiological phenotypes [1]. Epoxyeicosatrienoic acids (EETs), a major class of eicosanoids formed by Cytochrome P450 enzymes, play critical roles in endothelial migration, monocyte adhesion, tumor metastasis, and vasodilation, among other cell-type specific effects [2–4]. We recently showed that 11,12-EET enhances the specification of hematopoietic stem and progenitor cells (HSPCs) in developing zebrafish embryos, as well as the transplant of HSPCs in both fish and mice [5]. Despite the important physiological roles of EETs, their direct protein target(s) remains unknown.

Previous results indicate that EETs bind to at least one specific G-protein coupled receptor (GPCR). Nanomolar or even picomolar concentrations of 11,12 or 14,15-EET can elicit specific cellular phenotypes [6, 7], and a bead-tethered EET without the ability to cross the plasma membrane maintained its activity [8], suggesting the existence of a high-affinity, membrane-bound EET receptor. Chen *et al* (2011) demonstrated that U937 and other EET-responsive cell lines express a single high affinity EET receptor of about 47 kDa in size [9]. This is likely a GPCR, as EETs require G-protein signaling components to elicit many phenotypes [10–12]. For instance, we previously showed that EET’s enhancement of zebrafish hematopoiesis requires signaling via Gα12/13 [5].

As traditional biochemical methods have so far failed to identify EET receptors, here we used bioinformatic techniques to identify candidate receptors and assayed those candidates for EET-responsiveness *in vitro*. Only GPR132 (G2A), a previously described fatty acid receptor [13–16], showed responsiveness to EET. We demonstrated that GPR132 is required for EET-induced hematopoietic stem cell specification in the zebrafish, and for normal hematopoietic stem cell transplant in the mouse. Previously described fatty acid activators of GPR132 induced hematopoietic phenotypes in the zebrafish essentially identical to those observed with EET, further confirming that these molecules activate the same pathway. We performed structure-activity relationship analyses to determine the full range of GPR132 activators. Rather than a stereospecific, high-affinity receptor, our data show that GPR132 is likely a promiscuous receptor for a select panel of oxygenated polyunsaturated fatty acids, whose activity drives hematopoiesis in embryonic and adult vertebrate contexts.

## Results

### Identification of candidate EET receptors

EETs elicit phenotypes only in specific cell types, suggesting that a putative EET receptor might be selectively expressed in these cell types. We performed RNAseq profiling in duplicate on three EET-responsive human cell lines (U937 monocytes [17], EaHy endothelial cells [18], and PC3M-LN4 prostate cancer cells [4]), two of which show binding to a radiolabelled EET analog [9]. We profiled a fourth cell line (HEK293) that has no known responsiveness to EET and shows no such binding [9]. While we detected reads for hundreds of GPCRs (Fig. S1), only 37 GPCRs were expressed in common in all three EET-responsive cell lines above 0.3 fragments per kilobase per million reads (FPKM) (Fig. S2A), a conservative threshold for physiologically meaningful abundance. Of these, 27 were also expressed at moderate to high levels (FPKM>0.9) in our EET non-binding cell line (Fig. S2B). This left 10 candidate EET receptors that were expressed only in EET binding cell lines and missing from the non-binding cell line. All candidates had predicted molecular weights within 20% of the predicted 47 kDa size of an EET receptor [9], and candidates included both well-studied GPCRs such as the β-adrenergic receptor and prostaglandin receptors, as well as orphan GPCRs such as GPR132 and GPR135 (Fig. S2C).

### EET activates β-arrestin recruitment via GPR132 *in vitro*

GPCR activation causes recruitment of β-arrestin, which can be measured with the luminescence-based PathHunter assay [13, 15, 19]. We tested for EET-induced β-arrestin recruitment via each candidate GPCR, except GPR68, PTGER1 and LPAR6, which have no available PathHunter assays (Fig. 1). While candidates with known cognate small molecule ligands showed robust and dose-dependent activation by those ligands (Fig. 1C-F), EET showed no activity in these assays. CCRL2, a receptor for specific peptides, and GPR135, an orphan receptor, also failed to respond to EET (Fig. 1G,H). In contrast, 11,12-EET dose-dependently recruited β-arrestin to the GPR132 receptor, also known as G2A (Fig. 1B). Interestingly, two groups have previously assayed GPR132’s ability to respond to 14,15-EET and failed to see a high-affinity response [9, 20]. We found GPR132 to be more responsive to 11,12-EET than 14,15-EET (Fig. 1B). GPR132 is a member of the GPR4 family of GPCRs which are hypothesized to be lipid-sensing or acid-sensing receptors [21–24]. In addition to GPR132, this family includes GPR4, GPR65, and GPR68, an additional candidate from our bioinformatic analysis. No β-arrestin assay is available for GPR68, but 11,12-EET showed no activation of GPR4 or GPR65 (Fig. S3). Together, these results suggest 11,12-EET specifically binds to and activates GPR132, though this interaction may not be high-affinity.

**Figure 1:**
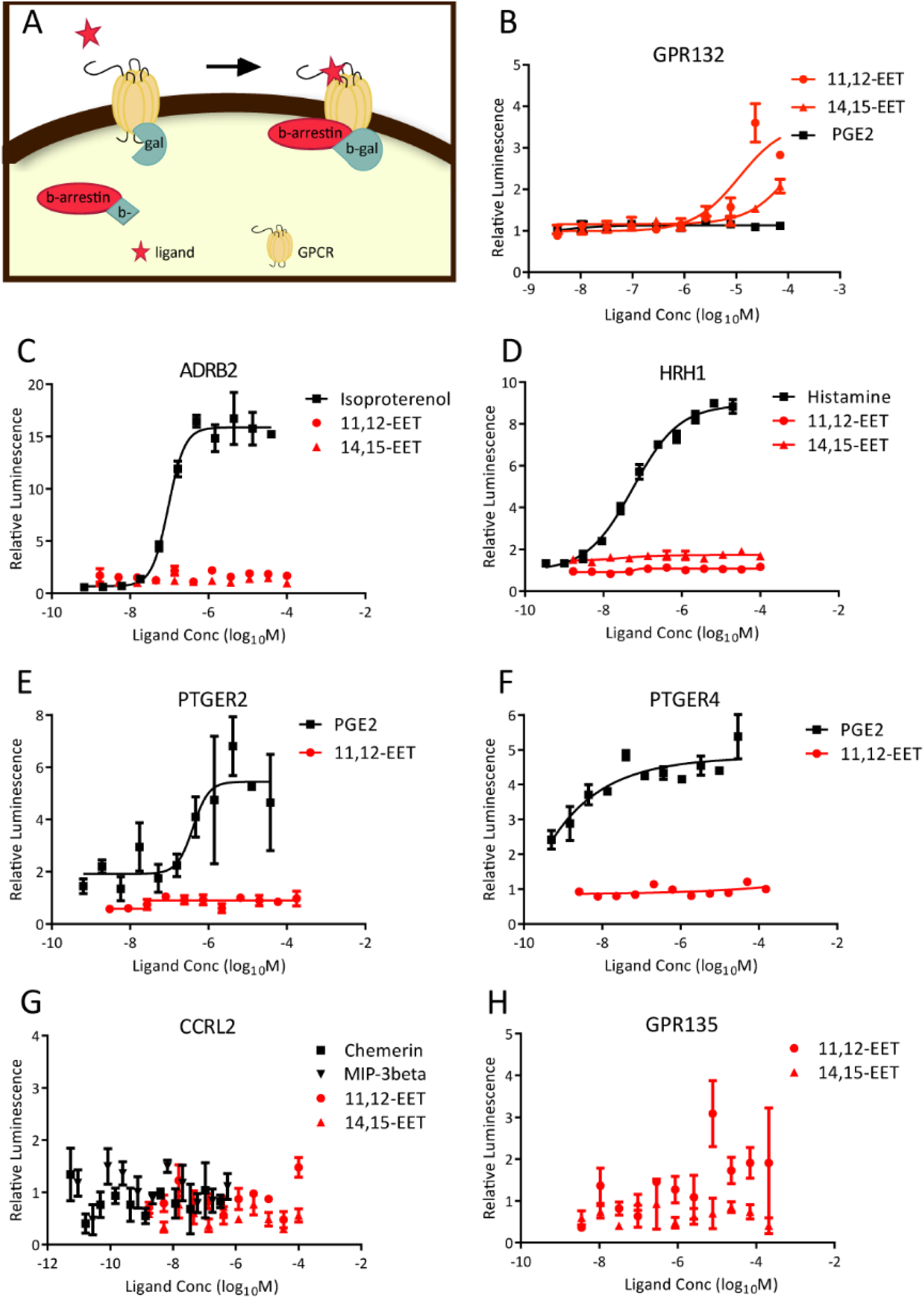
11,12-EET activates GPR132 *in vitro.* A) Diagram depicting the DiscoverX PathHunter β-arrestin assay. B-H) β-arrestin assays for candidate GPCRs treated with their known ligands or 11,12 or 14,15-EET. At each dose, cells were treated in triplicate and luminescence values were averaged. As basal luminescence varies significantly across assays for different proteins, luminescence values were normalized to the no treatment control. Each graph is representative of at least two similar experiments.

### *gpr132b* is required for EET-induced enhancement of zebrafish hematopoiesis

To determine whether GPR132 is required for EET phenotypes, we investigated EET-induced hematopoietic phenotypes in zebrafish embryos. We previously showed that EET treatment from 24-36 hours post fertilization (hpf) enhances specification of HSPCs in the aorta-gonad-mesonephros (AGM) region of the zebrafish trunk [5]. Later EET treatment enhances homing of HSPCs to the caudal hematopoietic territory (CHT), a developmental niche akin to the human fetal liver [5]. Zebrafish have two GPR132 homologs, *gpr132a* and *gpr132b*. At the amino acid level, they share 39% and 44% sequence similarity with human GPR132, respectively [22]. Previous RT-PCR studies showed *gpr132b* but not *gpr132a* expression in whole zebrafish embryos [25]. We were unable to expand a *gpr132a in situ* probe from embryonic RNA, perhaps due to low expression of this homolog. To determine where *gpr132b* is endogenously expressed in zebrafish embryos, we performed *in situ* hybridization in wildtype, untreated embryos at 24, 36, and 48hpf. *gpr132b* showed expression in the brain, including cells in the anterior lateral line nerve at all stages tested (Fig. S4). Staining was also seen in the posterior spinal chord at 24hpf (arrows). While *gpr132b* expression was not detectable by *in situ* hybridization in the AGM, it is expressed in the CHT. The CHT showed punctate expression (arrowheads) at all three stages, with strongest expression at 36hpf. This correlates with early stages of CHT colonization by myeloid precursors and HSPCs.

We designed a splice-blocking morpholino (MO) against *gpr132b* and injected 4-6 ng into 1-cell zebrafish embryos. Embryos were then treated from 24-36hpf with 5μM 11,12-EET and fixed at 36hpf to stain for *runx1*, a marker for HSPCs. Consistent with previous results, in uninjected or control MO-injected embryos, EET caused an increase in *runx1* staining in the AGM, and induced ectopic *runx1* staining in the tail mesoderm (Fig. 2) [5]. Injection of *gpr132b* MO completely blocked EET-upregulated *runx1* expression in the AGM, and partially blocked this upregulation in the tail (Fig. 2). This difference may reflect differing threshold requirements of *gpr132b* activity, with a higher level of *gpr132b* required in the AGM than the tail. Alternatively, the second zebrafish ortholog *gpr132a* may be able to compensate for the loss of *gpr132b* in the tail but not the AGM. Co-injections of human GPR132 RNA with the *gpr132b* MO restored the high AGM *runx1* staining, suggesting this MO effect was specific to loss of *gpr132b* (Fig. S5). GPR132 is thus required for EET’s enhancement of HSPC specification in zebrafish embryos.

**Figure 2:**
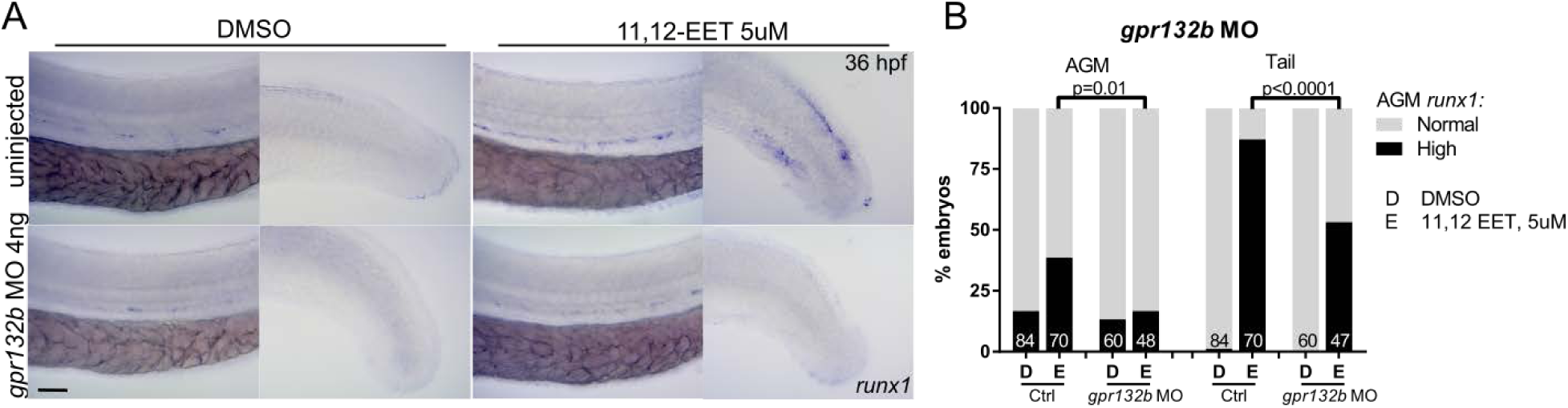
*gpr132b* is required for EET’s enhancement of zebrafish developmental hematopoiesis. A) Single-cell zebrafish embryos were injected with 4-6ng of *gpr132b* MO, control MO, or uninjected, then treated with DMSO or 5uM 11,12-EET beginning at 24hpf. Embryos were fixed at 36hpf and stained for *runx1* expression. B) Embryos were scored as having high or normal *runx1* expression in the AGM, and high (present) or normal (absent) *runx1* expression in the tail mesoderm. Graph presents summary of several experiments, total number of embryos at base of each column. Fisher’s exact test two-tailed. Ctrl includes control injected and uninjected embryos.

### GPR132 knockout mice show impaired long-term HSC transplant

GPR132 KO mice have normal overall physiology, although they are known to show an age-related autoimmune disease featuring an increase in B-cells and T-cells, and lymphocytic infiltration of tissues [26]. We found that 3.5 - 5 month old GPR132 KO mice have normal distributions of B-cells, T-cells, and granulocytes, as well as comparable numbers of phenotypic long-term HSCs, short-term HSCs and multipotent progenitors compared to their wildtype or heterozygous siblings (Fig. S6). To stringently test HSPC function, we performed limiting dilution whole marrow transplants from GPR132 knockout and heterozygous mice. We transplanted donor cells from CD45.1+, GPR132 -/- or +/-mice together with wildtype CD45.2 competitor cells into irradiated CD45.2 recipients and observed peripheral blood chimerism over time (Fig. 3). At a limiting, 10,000 cell dose, GPR132 knockout marrow shows a functional defect in competitive long-term engraftment compared to sibling heterozygous marrow, indicating that GPR132 is required for normal HSC function. This defect was consistent in all blood lineages (Fig. S7A,B) and in the marrow (Fig. S7C), and resulted in a significantly smaller proportion of knockout mice displaying multi-lineage chimerism (Fig. S7D). No difference was seen between heterozygous and knockout marrow at the higher cell dose. There was no difference in lineage contribution between heterozygous or knockout marrow at either cell dose (Fig. S7E). We chose representative recipient mice from each cell dose and used marrow from these to perform secondary transplants (Fig. 3, Fig. S7). Generally, peripheral blood and marrow chimerism correlated between primary and secondary recipients (Fig. S8), and knockout marrow continued to show impaired engraftment at the limiting, 10,000 cell dose (Fig. 3, Fig. S7A,B). Overall, this transplant data suggests that GPR132 knockout mice have a decreased number of functional long-term HSCs compared to their heterozygous siblings, even in the absence of exogenous EET stimulation. Therefore, it would be difficult to interpret the effects of exogenous EET stimulation in these transplants. The question of how GPR132 signals endogenously in mammalian hematopoietic niches, which could be constitutive, or in response to endogenous ligands, is an important area for further study.

**Figure 3:**
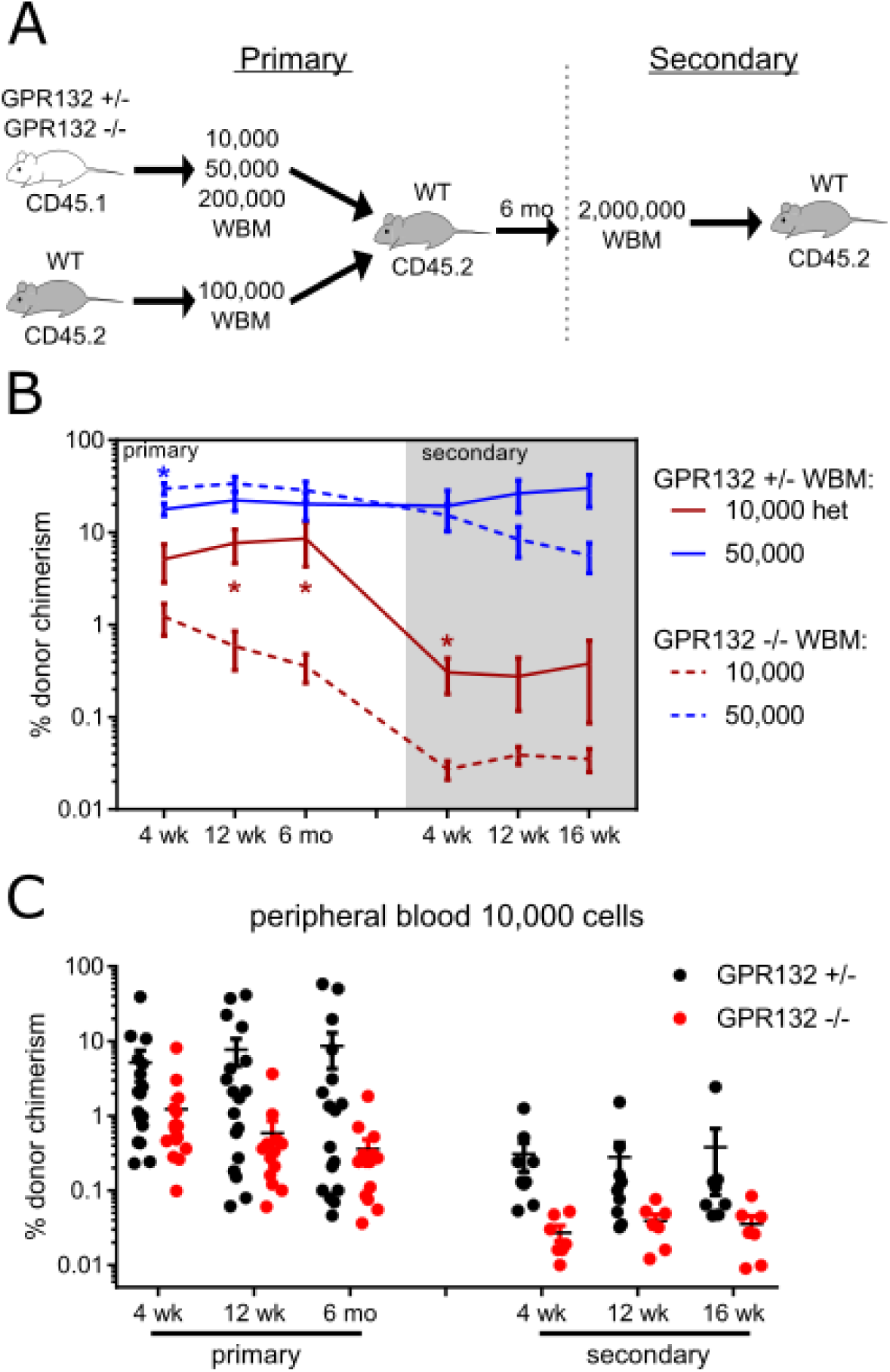
GPR132 is required for normal marrow transplant in the mouse. A) 10,000-500,000 donor cells were combined with 100,000 CD45.2 wildtype competitor cells and transplanted into lethally irradiated CD45.2 recipients. Mice were bled at indicated time-points and cells were stained for donor chimerism. At 6 months post-transplant, representative primary recipients were sacrificed and 2 million whole marrow cells were transplanted into secondary recipients. B) Peripheral blood chimerism in transplant recipients. Graph summarizes data from three separate experiments with a total of 14-17 primary recipients per condition. 3-6 primary recipients were used as donors for 3 secondary recipients each. Mean with SEM is plotted. *, p<0.05, student’s T-test. C) Peripheral blood contribution of 10,000 donor cells over time for each genotype in primary and secondary transplant, individual mice shown.

### Known GPR132 agonists activate hematopoiesis in developing zebrafish

GPR132 is a reported receptor for bioactive lipids including 9-HODE and 11-HETE [13–16]. Like 11,12-EET, these molecules are oxygenated, polyunsaturated free fatty acids (Table 1). We confirmed these previous data and showed that both of these molecules recruited β-arrestin downstream of GPR132 *in vitro* dose-dependently (Fig. 4A). Additionally, we treated developing zebrafish embryos with 9-HODE and 11-HETE from 24-36hpf, then fixed the embryos and performed an *in situ* hybridization to examine expression of the hematopoietic markers *runx1* and *c-myb*. 9-HODE and 11-HETE produced EET-like phenotypes in zebrafish embryos, increasing *runx1/c-myb* expression in both the AGM and tail of the fish (Fig. 5, Table 1). The tail staining of *runx1/c-myb* is a phenotype we have uniquely seen with EET treatment, despite conducting a wide variety of chemical treatments of embryos in our lab. The high similarity between EET, 9-HODE, and 11-HETE phenotypes strongly suggests these molecules are activating the same receptor, namely GPR132.

**Figure 4:**
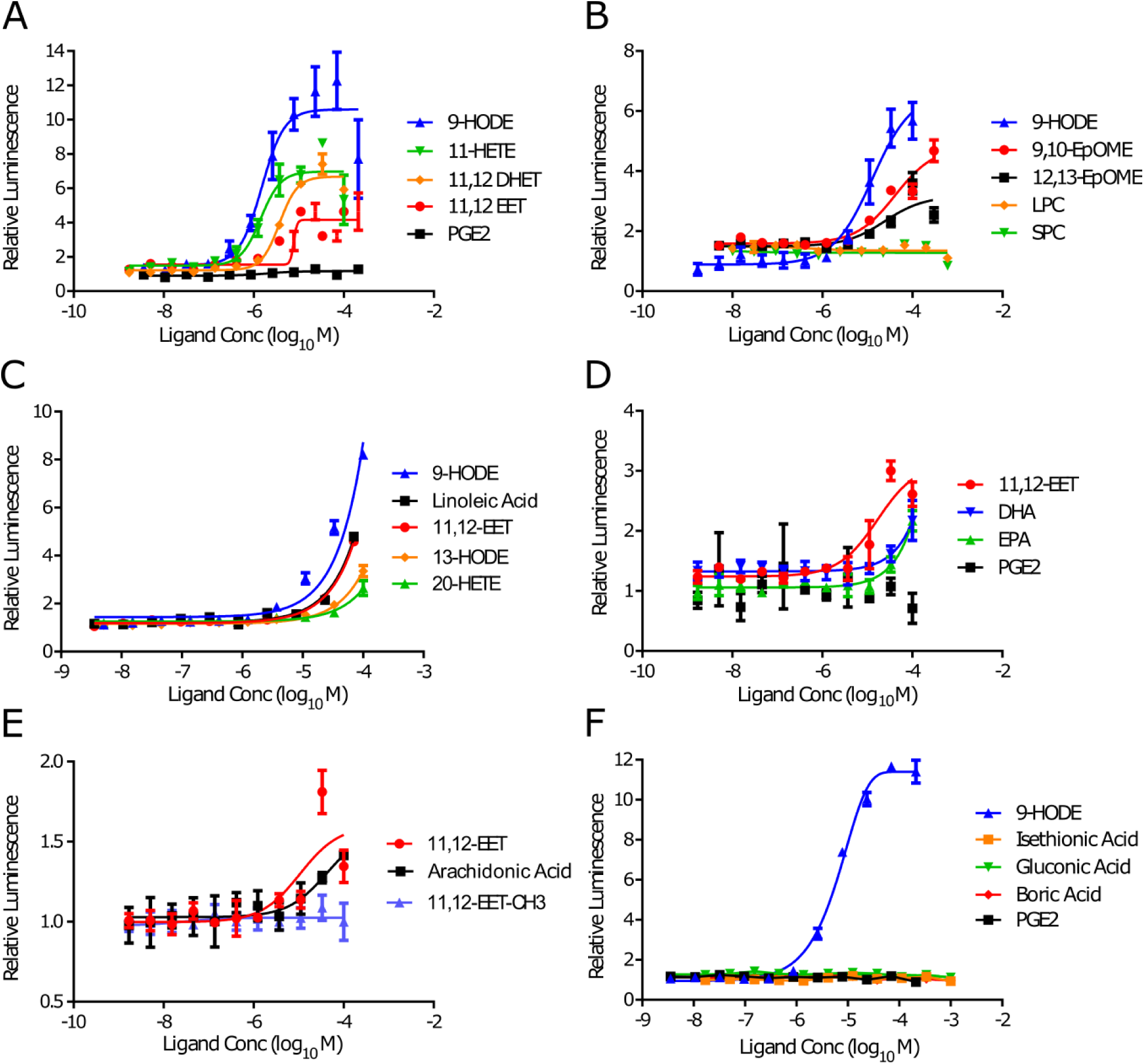
Diverse free fatty acids activate GPR132 signaling *in vitro*. A-F) GPR132 β-arrestin recruitment assays. Small molecules were treated in triplicate and each graph represents at least two similar experiments. Error bars represent SEM. Note differing y-axis scales.

**Figure 5:**
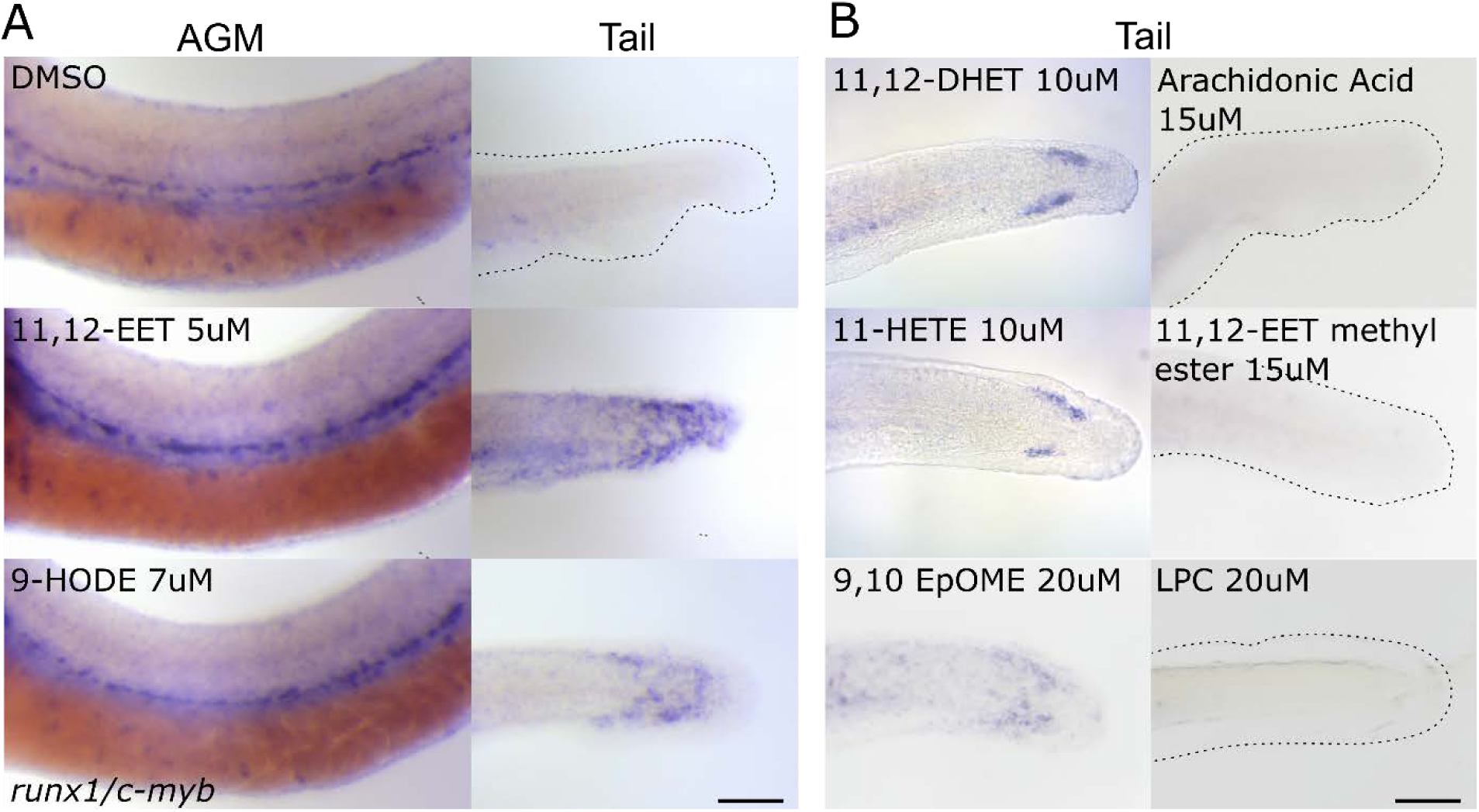
Fatty acid GPR132 activators enhance expression of HSPC markers in zebrafish embryos. A) *in situ* hybridization of zebrafish embryos treated with DMSO, 11,12-EET, or 9-HODE showing AGM and tail expression of runx1/c-myb. For full quantification, see Table 1. B) *in situ* hybridization of zebrafish embryos treated with diverse free fatty acids, EET-methyl ester, and lysophosphatidylcholine (LPC) showing tail expression of runx1/c-myb. For full quantification, see Table 1. Embryos were dechorionated at 24hpf and incubated with the indicated quantities of small molecule from 24-36hpf. Embryos were fixed at 36hpf and stained using a probe combination of runx1 and c-myb. Scale bar is 100μm.

**Table 1.** Diverse free fatty acids enhance HSPC specification in zebrafish embryos. Embryos were treated with the indicated concentrations of drug from 24-36hpf, then fixed and stained for *runx1/c-myb* expression. AGMs were scored as having high, medium, or low expression, and tails were scored as having high (present) or low (absent) expression. Doses which caused morphological defects or circulation problems were disregarded. Data here is a summary of many experiments. Percentages of embryos having high expression in the AGM and tail are shown, as well as the total number of embryos assayed for each molecule. Each drug was tested in at least two separate experiments. Each experiment included clutch-matched negative (DMSO or ethanol) and positive (11,12-EET) control treatment conditions.

| Drug | Structure | Dose<br>(μM) | AGM | | Tail | |
| --- | --- | --- | --- | --- | --- | --- |
|  |  |  | % High | N | % High | N |
| DMSO |  |  | 12.6 % | 388 | 0.5 % | 393 |
| Ethanol |  |  | 12.2 % | 278 | 0.0 % | 278 |
| 11,12-EET |  | 5 | 41.2 % | 585 | 63.8 % | 585 |
| 11,12-DHET |  | 3 | 24.2 % | 33 | 3.0 % | 33 |
|  |  | 5 | 51.6 % | 31 | 16.1 % | 31 |
|  |  | 10 | 61.0 % | 77 | 48.1 % | 77 |
| 9-HODE |  | 7 | 17.2 % | 29 | 48.4 % | 31 |
|  |  | 10 | 57.4 % | 47 | 10.6 % | 47 |
| 13-HODE |  | 5 | 23.1 % | 26 | 0.0 % | 27 |
|  |  | 8 | 6.3 % | 32 | 3.3 % | 30 |
|  |  | 12 | 14.9 % | 67 | 0.0 % | 66 |
|  |  | 20 | 17.2 % | 29 | 0.0 % | 29 |
| 11-HETE |  | 3 | 14.3 % | 28 | 7.1 % | 28 |
|  |  | 5 | 15.2 % | 33 | 12.1 % | 33 |
|  |  | 10 | 68.1 % | 69 | 13.0 % | 69 |
| 20-HETE |  | 5 | 19.0 % | 21 | 50.0 % | 22 |
|  |  | 8 | 13.3 % | 30 | 26.7 % | 30 |
|  |  | 12 | 13.0 % | 23 | 72.7 % | 22 |
| 9,10-EpOME |  | 5 | 14.0 % | 57 | 0.0 % | 57 |
|  |  | 10 | 37.0 % | 54 | 13.0 % | 54 |
|  |  | 20 | 32.8 % | 58 | 81.0 % | 58 |
| 12,13-EpOME |  | 5 | 16.7 % | 42 | 0.0 % | 42 |
|  |  | 10 | 31.1 % | 61 | 1.7 % | 60 |
|  |  | 20 | 20.4 % | 49 | 0.0 % | 49 |
| Arachidonic Acid |  | 10 | 4.8 % | 42 | 0.0 % | 42 |
|  |  | 15 | 17.4 % | 23 | 0.0 % | 23 |
| Linoleic Acid |  | 5 | 12.0 % | 25 | 0.0 % | 26 |
|  |  | 8 | 8.5 % | 59 | 0.0 % | 60 |
|  |  | 12 | 12.5 % | 56 | 0.0 % | 54 |
|  |  | 20 | 29.2 % | 24 | 4.2 % | 24 |
| Eicosapentaenoic Acid |  | 7 | 8.8 % | 34 | 0.0 % | 34 |
|  |  | 15 | 34.2 % | 38 | 0.0 % | 38 |
|  |  | 30 | 20.5 % | 44 | 10.0 % | 40 |
| Docosahexaenoic Acid |  | 7 | 3.4 % | 59 | 0.0 % | 59 |
|  |  | 15 | 13.7 % | 51 | 0.0 % | 51 |
|  |  | 30 | 11.1 % | 54 | 0.0 % | 54 |
| 11,12-EET-methyl ester |  | 5 | 22.2 % | 27 | 0.0 % | 27 |
|  |  | 7 | 23.2 % | 82 | 0.0 % | 82 |
|  |  | 10 | 17.5 % | 57 | 0.0 % | 57 |
|  |  | 15 | 15.1 % | 53 | 1.9 % | 53 |
| LPC |  | 10 | 35.3 % | 34 | 0.0 % | 34 |
|  |  | 20 | 35.7 % | 28 | 0.0 % | 28 |
| SPC |  | 10 | 24.3 % | 37 | 0.0 % | 37 |
|  |  | 20 | 17.6 % | 17 | 0.0 % | 17 |

### GPR132 is responsive to a range of oxygenated fatty acids

As GPR132 showed comparable reactivity to 11,12-EET, 9-HODE, and 11-HETE, we sought to further explore potential GPR132 ligands by conducting structure-activity relationship analyses. We assayed a variety of bioactive lipids for their ability to recruit β-arrestin to GPR132 *in vitro* and to enhance *runx1/c-myb* expression in zebrafish embryos *in vivo*. We found that simple chain, oxygenated, unsaturated free fatty acids including 11,12-EET, 11-HETE, 9-HODE, and 11,12-DHET robustly enhanced both of these phenotypes (Table 1, Fig. 4A, Fig. 5). More saturated lipids such as 9,10 and 12,13-EpOME showed lower affinity in β-arrestin assays (Fig. 4B) and showed little activity in the zebrafish *in vivo* (Fig. 5B, Table 1). Fatty acids such as 13-HODE and 20-HETE, with their oxygenated groups displaced farther from the carboxylic acid, similarly showed decreased activity *in vitro* (Fig. 4C) and *in vivo* (Table 1). Oxygenation seems to contribute to GPR132 activation, as the unoxygenated free fatty acids linoleic acid, eicosapentaenoic acid, docosahexaenoic acid, and the unoxygenated EET precursor arachidonic acid showed very little activity *in vivo* (Table 1, Fig. 5B). *In vitro* these molecules also showed reduced efficacy compared to 11,12-EET (Fig. 4D,E), with the exception of linoleic acid, which behaved equivalently to 11,12-EET in the β-arrestin assay (Fig. 4C). 11,12-EET-methyl ester, which replaces the carboxylic acid of 11,12-EET with a methyl group, had no activity either in β-arrestin assays or in zebrafish embryos, suggesting the carboxylic acid is absolutely required for binding to GPR132 (Table 1, Fig. 4E, Fig. 5).

Previous studies have hypothesized that GPR132 and its near family members function as acid-sensing receptors [21–24, 27], or as receptors for lysophosphatidylcholine (LPC) and sphingosylphosphorylcholine (SPC) (reviewed in [28]). These results have been controversial, and direct binding experiments linking LPC to GPR132 were later retracted [29, 30]. We therefore assayed the abilities of these molecules to activate GPR132. In our hands, lactic acid, LPC, and SPC all failed to induce β-arrestin recruitment by GPR132 (Fig. 4B, Fig. S9) and failed to enhance HSPC marker staining in zebrafish (Fig. 5B, Fig. S9). Three additional organic acids also did not cause recruitment of β-arrestin by GPR132 (Fig. 4F). Collectively, our data point to GPR132 as a receptor for medium-chain, oxygenated free fatty acids, rather than a broadly pH-sensitive receptor, or an LPC/SPC-responsive receptor.

## Discussion

We employed a bioinformatic pipeline and experiments in zebrafish, a genetically and chemically tractable model organism, to address the long-standing question of the identity of the EET receptor. We found that 11,12-EET enhances HSPC specification in zebrafish embryos by activating the receptor GPR132. This receptor is responsive to a select panel of oxygenated polyunsaturated fatty acids, suggesting that it can integrate these diverse signals *in vivo* to regulate hematopoiesis. In addition, GPR132 appears to be a conserved regulator of HSPC function in mammals, where loss of this receptor leads to impaired long-term reconstitution after primary and secondary transplant.

In contrast to our assays, classically-studied EET phenotypes often display high affinity and high stereospecificity (reviewed in [2]). For instance, EET stimulation of BK(Ca) channels is a high affinity response [7] that is specific to EET and not its metabolite DHET [31]. However, in our β-arrestin assays and zebrafish embryo treatments, EET and DHET were both active, along with additional polyunsaturated, oxygenated fatty acids. Accurate affinity measurements cannot be made from either of our methods due to tissue accessibility challenges in the zebrafish and serum interference with binding in β-arrestin assays [32]. Direct biochemical interrogation of EET-GPR132 binding is still needed. We attempted to assay the ability of 11,12-EET to compete with a tritium-labelled 9-HODE (synthesized by American Radiolabeled Chemicals) for binding to GPR132. However, we were only able to obtain H3-9-HODE with a very low specific activity (0.13 Ci/mMol), which impeded an accurate assessment of binding. Other proposed EET receptors include TRPV/TRPC receptors [33–35], PPARs [36], prostaglandin receptors [20, 37], and FFAR1 (also called GPR40) [38]. We do not see expression of FFARs in zebrafish embryos (data not shown), and saw no response of two prostaglandin receptors to EET *in vitro* (Fig. 1G,H). Voltage gated channels or nuclear hormone receptors do not explain the G-protein dependence of EET phenotypes. Interestingly, though, 9-HODE and 13-HODE are also reported to activate TRPV1 [39]. Our results point to GPR132 as a potentially low-affinity EET and oxygenated fatty acid receptor with physiological relevance in hematopoiesis. We cannot rule out the existence of additional EET receptors, responsible for other EET phenotypes, which may have high-affinity, stereospecific interactions.

GPR132 itself is controversial molecule which has been described variously as a receptor for free fatty acids [13–15], a receptor for LPC and SPC, or a proton and pH sensitive receptor (reviewed in [40, 41]). We saw no evidence of LPC-, SPC-, or pH-responsiveness of GPR132. Our data instead support the role of GPR132 as a free fatty acid receptor, and further define that maximum activity is achieved with medium chain, multiply unsaturated, oxygenated fatty acids. Hetero-dimerization of GPR132 with other GPCRs could explain the differing ligand sensitivities seen by many groups. One overexpression study suggested that GPR132 hetero-dimerizes with its close family member, GPR68 [42], which was an additional candidate EET receptor in our bioinformatic analysis. Hetero-dimerization with GPR68 or other GPCRs could affect GPR132’s acid responsiveness, or its affinity or stereospecificity for fatty acids.

GPR132 is expressed in macrophages, and its previously described physiological roles include regulating macrophage activation [27, 43] and, in aging mice, the production of lymphoid cells [26]. We have described for the first time a role for GPR132 in regulating HSPCs. This regulation could function autonomously in HSPCs, or could occur via macrophages, which are present in developmental and adult HSPC niches and are known to regulate HSPC function (reviewed in [44]).

Beginning with a novel use of bioinformatic analysis to de-orphan the bioactive lipid 11,12-EET, we have uncovered a fatty acid-GPR132 signaling axis which regulates developmental and adult hematopoiesis. These results add to prior studies by our group demonstrating a role for prostaglandins in regulating HSPC development and transplant [45], as well as earlier work exploring the roles of lipid mediators such as HETEs, HODEs, and leukotrienes in regulating inflammation [46–49], as well as B-cell production, adhesion, and Ig production [50–53], and T-cell function [54–56]. Together, these reports show that complex networks of lipid signaling molecules regulate diverse biological phenomena, including hematopoiesis and stem cell dynamics [57]. GPR132 may represent a node in this network, able to integrate the input of diverse lipid-derived signals into a single pro-hematopoietic output. Systematic understanding of these networks is sorely needed.

## Methods

### Materials

All fatty acids, EET-methyl ester, and isoproterenol were purchased from Cayman Chemicals at 5-10mM stock concentrations in ethanol or DMSO. All chiral molecules were purchased as racemic mixtures. Histamine dihydrochloride (H7250), LPC (L1881), SPC (S4257) and lactic acid (L6661) were purchased from Sigma-Aldrich and diluted in ethanol. Recombinant human chemerin and MIP-3β were purchased from PeproTech, Inc.

### PathHunter β-arrestin assays

PathHunter β-arrestin assays were purchased from DiscoverX and performed in 384 well plates according to manufacturer instructions. Cells were thawed and plated in Cell Plating 0 buffer, which contains 1% heat-inactivated FBS (ADRB2, PTGER4, GPR4), Cell Plating 1 buffer, which contains 2% charcoal/dextran treated serum (GPR132, CCRL2, GPR135, GPR65), or Cell Plating 2 buffer, which contains 10% heat-inactivated FBS (HRH1, PTGER2,). In each experiment, small molecules were plated in triplicate across a range of concentrations and incubated with reporter cells for 90 minutes at 37°C. Luminescence was developed using DiscoverX reagents, and luminescence values were normalized to baseline within each experiment. Each small molecule was tested in a minimum of two separate experiments. Small molecule stocks were prepared in DMSO or ethanol. EC50 values and non-linear regressions were performed in GraphPad Prism using the log(agonist) vs response function.

### Zebrafish embryo experiments

Zebrafish were housed and cared for according to IACUC protocol 14-10-2789. Casper or AB zebrafish were used for all experiments. For morpholino injections, embryos were injected at the 1 cell stage with 4-6ng MO, mixed with phenol red as an indicator of injection success. The Gpr132b_sp MO was produced by GeneTools, with a sequence of TAAAATGGCGTTGCTCTTACCTCTA. The Standard Control morpholino was also obtained from GeneTools, with the sequence CCTCTTACCTCAGTTACAATTTATA. For small molecule treatments, embryos were dechorionated at 24hpf, and small molecule was added to E3 media and embryos in 12-well plates. Embryos were fixed at 36hpf in 4% PFA, and *in situs* were performed as described [58].

### Mouse transplant

Mice were housed and cared for according to IACUC protocol 15-06-2964. GPR132 knockout mice (Jackson Labs #008576) were kindly provided by Donna Bratton. Mice were genotyped for GPR132 status by PCR as previously described [26]. Donor and competitor marrow from femur, tibia, and hip was harvested by crushing. The indicated number of donor cells were mixed with 100,000 CD45.2 wildtype (Jackson Labs #000664), sex- and age-matched competitor marrow cells. Cells were transplanted retro-orbitally into sex- and age-matched CD45.2 wildtype recipients, which had received 10 gy irradiation in a split dose. At 6 months post-transplant, recipient mice were sacrificed and their marrow was analyzed for CD45.1 chimerism and used in secondary transplants. In secondary transplants, 2 million cells from a total of 24 primary recipients were transplanted into 3 CD45.2 secondary recipients each.

### Antibody staining and flow cytometry analysis

Mouse peripheral blood was incubated with ACK lysing buffer (Thermo Fisher) in order to deplete red blood cells. Lysing buffer was washed off, and the remaining cells were incubated for 1 hour at 4°C with 1:100 dilutions of the following antibodies: Ter119-PE-Cy5 (red blood cells, eBioscience), CD3-APC (T-cells, eBioscience), B220-Pacific Blue (B-cells, eBioscience), Gr1-PE-Cy7 (clone RB6-8C5, granulocytes, eBioscience), CD45.1-PE (BD Pharmingen), and CD45.2-FITC (BD Pharmingen). All stainings were performed in PBS at 4°C for 1 hour. Antibodies were then washed off and cells were analyzed on an LSRII Flow Cytometer. FlowJo software was used to process the data.

## Acknowledgements

We thank E. Fast for valuable discussion. This work was supported by NIH grants R01HL04880, P01HL032262, U01HL100001, R24DK092760, and U01HL134812 (to LIZ), 5F31HL129517-02 (to JLL), and P01GM095467 (to CNS).

